# Selective reduction of *KCNA4* in vulnerable glutamatergic-serotonin neurons of the dorsal raphe nucleus in Alzheimer’s Disease

**DOI:** 10.1101/2025.10.17.683113

**Authors:** Louis J. Kolling, Nagalakshmi Balasubramanian, Shafa Ismail, Alexander J. Feller, Jessica Marie Hunter Alberhasky, Ruixiang Wang, Luke Jennings, Marco Hefti, Catherine Anne Marcinkiewcz

**Affiliations:** Department of Neuroscience and Pharmacology, University of Iowa, Iowa City, Iowa, USA; Department of Cellular and Systems Pharmacology, University of Florida, Gainesville, USA; Department of Pathology, University of Iowa, Iowa City, Iowa, USA

**Keywords:** Tau, Dorsal raphe nucleus, spatial transcriptomics, 5HT/glut neurons, Alzheimer’s disease

## Abstract

**INTRODUCTION:** We previously demonstrated that htau mice recapitulate many neuropsychiatric features of early Alzheimer’s disease (AD), and that the dorsal raphe nucleus (DRN) contains distinct subregions. Herein, we investigate vulnerability of the centromedial DRN to pathologically-phosphorylated tau (pTau), a region composed predominantly of dually serotonergic/glutamatergic (5HT/glut) neurons.

**METHODS:** We use computational, molecular, biophysical, and behavioral techniques to assess the centromedial DRN across preclinical and post-mortem settings.

**RESULTS:** The centromedial DRN contains 5HT/glut neurons that differentially express ion-channel genes in the htau mouse. 5HT/glut neurons exhibit increased excitability, which we demonstrate may dually promote pTau accumulation and the severity of depressive-like behaviors in htau mice. At Braak 2, *KCNA4* is reduced in 5HT/glut neurons in AD, which are especially vulnerable to pTau compared to 5HT-nonglut neurons.

**DISCUSSION:** Tau-mediated dysfunction of the DRN may be driven by changes in ion channel activity that concomitantly promote the spread of pTau in Braak progression.

## 1 INTRODUCTION

Alzheimer’s disease (AD) is the fifth-leading cause of death in older adults [1] and there are currently no effective therapeutics to halt, reverse, or prevent its development. This is due in part to the complex clinical presentation of AD, a mixed proteinopathy that often co-presents with other disease [2–6]. Of these various factors, a hyper-phosphorylated form of the tau protein (pTau) is heavily implicated in the earliest AD-related cognitive decline [7–13]. Growing evidence suggests that pTau first spreads from areas of the brainstem [14–17], including the serotonin-producing (5-HT) raphe nuclei. Dysfunction of 5-HT circuitry may drive many of the neuropsychiatric symptoms that develop during the prodromal phase of AD and precede the onset of cognitive decline [18], including depression, anxiety, and disruptions to sleep architecture [19–22]. Therefore, the 5-HT system represents a critical therapeutic target to abrogate the spread of tau pathology in AD.

We previously found that mice that over-express non-mutant human tau (htau mice) recapitulate many of the neuropsychiatric features of early AD, and we observed subtle changes in the action potential (AP) kinetics and innate biophysical properties of 5-HT neurons in these mice [23]. These characteristics, which are primarily governed by ion channels, may play a role in the dysregulation of neuronal excitability that is common in Alzheimer’s disease and various other neurodegenerative diseases [24]. We then used the Visium spatial gene expression platform to profile the htau dorsal raphe nucleus (DRN), the largest of the serotonin-producing nuclei [25]. We characterized three computationally-distinct subregions of the DRN—the centromedial, ventrolateral, and dorsolateral DRN. Within the centromedial (CM) DRN, we identified differential expression of ion-channel genes, which may be crucial for the regulation of 5-HT neuron excitability and action. Pathologically phosphorylated tau has previously been shown to disrupt neuronal excitability [26], which may concomitantly exacerbate the trans-synaptic spread of pTau [27,28].

The goal of this study was to use our new understanding of DRN subregions to enhance the preclinical-to-clinical translatability of the htau mouse model. Our choice of the CM DRN was inspired, in part, by our previous identification of a regional “marker gene” that could possibly be used to directly test our preclinical findings in post-mortem human DRN tissue from patients who had been diagnosed with AD. We hypothesized that the CM DRN would primarily contain 5-HT neurons that are dually glutamatergic (5HT/glut neurons), and that 5HT/glut neurons of htau mice would recapitulate the genetic and pathological features of human 5HT/glut neurons in AD—enabling the continued interrogation of AD etiology in the htau mouse model. Here, we report differential gene expression of ion channels, such as *Kcna4* (Kv1.4) and *Slc24a5* (NCKX5) in htau 5HT/glut neurons that accompanies an increase in their evoked excitability and action-potential df/dt (rate of change of frequency). We further demonstrate that chronic hyper-excitability may affect long-term pTau accumulation in htau mice, independent of cell type. Following post-mortem human AD investigation, we find that 5HT/glut neurons differentially express *KCNA4* in AD, and that 5HT/glut neurons are especially vulnerable to tau pathology as compared to 5HT-nonglut neurons (non-glutamatergic 5-HT neurons). Modulation of Kv1.4 activity and gene expression in 5HT/glut neurons may represent a critical therapeutic target for the treatment of tau-based AD.

## 2 MATERIALS AND METHODS

### 2.1 Ethical approval

All animal procedures were reviewed and approved by the University of Iowa (UI) Office of Animal Resources (Protocol 4032080) that abided by the AVMA and NIH.

### 2.2 Animals

Male C57 mice (C57/BL/6J mice; catalog #000664, Jackson Labs, Bar Harbor, ME, USA; RRID:IMSR_JAX:000664) and htau hemizygous mice (catalog #005491, Jackson Labs, Bar Harbor, ME, USA; RRID:IMSR_JAX:005491), were purchased for use. 5 C57 and 5 htau hemizygous mice were aged to 16 weeks and used for RNAscope. 8 C57 and 8 htau hemizygous mice were aged to 16 weeks and used for electrophysiology. 37 htau hemizygous mice underwent DREADDs surgery at 8 weeks’ age and were aged to 16 weeks for behavioral and pathological assessments. Since only male mice were used for the Visium Spatial Gene Expression experiment, only male mice were used for these subsequent experiments. 13 C57 mice and 50 htau mice were used for experiments (63 mice total).

All mice were housed in a temperature- and humidity-controlled AALAC-approved vivarium at UI on a standard 12 h/12 h dark/light (reverse) cycle in accordance with institutional requirements. Mice were housed in conventional-style rodent cages with shredded paper bedding, containing separate food and water that could be obtained *ad libitum*. Mice were maintained on chow that was composed of 14% kcal fat, 60% kcal carbohydrate, and 26% kcal protein (Catalog #5P76; Land O’Lakes, Arden Hills, MN, USA).

### 2.3 Justification of sex

Given the complex interplay between estrogens and 5-HT neurons of the DRN, the incorporation of female mice, and consequently, female AD tissue, is beyond the scope of this report. 5-HT neurons of the DRN differentially express estrogen receptor beta (ERβ) in a sex-dependent manner [29,30]. Activation of ERβ directly regulates the expression of *Tph2* [31], the primary rate-limiting enzyme for 5-HT production, directly confounding any assessments of 5-HT neuronal function, activity, and related behaviors. Additionally, estrogens affect the expression of monoamine oxidase B, an enzyme responsible for 5-HT degradation, in a region- and sex-specific manner [32]. The current study will provide a point of comparison for a future study that focuses on the relationship between tau pathology and 5-HT in females.

### 2.4 Visium spatial transcriptomics

Cluster assignment and identification of DEGs have been described in detail previously [25]. Advaita Bio’s iPathwayGuide [33,34] was used to identify relevant cellular pathways associated with DEGs, particularly those associated with disease-related processes. Identification of candidate “marker genes” for DRN subregions was then performed using the differential expression tool within the Loupe Browser (v 7.0). The full dataset of centromedial DRN DEGs was then compared with the Agora human AD database.

### 2.5 DREADDs experiment

#### 2.5.1 DREADDs surgeries

37 male htau mice underwent surgery at 8 weeks’ age. All stereotactic infusions targeted the CM DRN using the following coordinates (relative to the bregma, in mm): AP -4.6, ML 0, DV -3.6. Virus titers were approximately 2.5 x 10^13^, and each mouse received a single 300 nl infusion. 9 “control” mice received AAV8-hsyn-mCherry, 14 mice received AAV8-hsyn-hM3Dq-mCherry, and 14 mice received AAV8-hsyn-hM4Di-mCherry. All surgeries were completed within a 3-day period.

#### 2.5.2 Chronic CNO challenge

After a 7-day recovery period, mice were continuously administered the DREADD agonist CNO DHC (clozapine-n-oxide dihydrochloride, Hello Bio #HB6149, Princeton, NJ, USA) at a concentration of 6.06 mg/ml (equivalent to 5 mg/ml CNO) via water bottle in a home-cage environment. Fresh CNO solution was prepared thrice-weekly in tap water, and water bottles were wrapped in foil to shield against UV degradation. Mice were continuously administered CNO for a period of 6 weeks and were regularly assessed for water consumption (∼4ml/day/mouse). At the end of the 6-week CNO challenge, 2 mice from each DREADDs group were sacrificed for *Fos* assessment to determine the ability of DREADDs to chronically modulate the transduced neurons (**Section 2.6.2**).

#### 2.5.3 DREADDs behavior assessment

After the 6-week CNO challenge, mice were administered normal drinking water for an additional 7 days to “wash out” the effects of the CNO as a DREADDs ligand. This was performed to prevent residual chemogenetics effects from confounding the behavioral assessments. At the end of the washout period, 2 mice from each DREADDs group were sacrificed for *Fos* assessment to determine whether CNO washout was successful (**Section 2.6.2**). Mice underwent behavioral assessments in the following order: novelty-induced suppression of feeding, novel-object recognition, elevated plus maze, social interaction test. No rest days were provided between tests to shorten the period between CNO challenge and histological assessment of pTau.

##### 2.5.3.1 Novelty-induced suppression of feeding

48 hours before testing, mice were each given a single piece of Froot Loops cereal (Kellogg’s) in their home cage. 24 hours before testing, all chow was removed from the home cage, and mice remained fasted until testing was complete. On test day, mice were transferred to a testing room and allowed to acclimate for 1 hour. Mice were then individually placed into an open arena (50 x 50 x 25 cm; 20–25 lux) that contained a single Froot Loop in the center of the arena. Mice were allowed to explore the arena for 10 mins while the tester was absent from the room, and latency to feed on the Froot Loop was measured *post hoc* using recorded video. Mice were then returned to their home cage and allowed to feed freely on the Froot Loop. Any mouse that did not feed on the Froot Loop in their home cage after the test was excluded from data collection.

##### 2.5.3.2 Novel-object recognition

Short-term memory was assessed using the novel object recognition (NOR) test as described previously [35]. The test consisted of three phases: habituation, acquisition, and testing phase. During habituation (day 1), mice were allowed to freely explore the empty arena for 10 min. After 24 hours, in the acquisition (training) phase, each mouse was placed in the same arena containing two identical objects positioned equidistantly and allowed to explore for 10 min. Two hours later, during the test phase, one of the familiar objects was replaced with a novel object, and the mouse was allowed to explore both objects for 5 minutes. Exploration was defined as the mouse directing its nose toward, sniffing, or touching the object. Exploration time was recorded by an observer blinded to the experimental groups. The discrimination index (DI) was calculated as (N − F) / (N + F), where *N* represents the time spent exploring the novel object and *F* represents the time spent exploring the familiar object. A positive DI indicates preference for the novel object, whereas a negative DI indicates preference for the familiar object. Percent novel preference was calculated as *(N / (N + F)) × 100*. Reduced discrimination and diminished novel object preference during the test session indicate impairments in recognition memory.

##### 2.5.3.3 Elevated plus maze

Anxiety-like behavior was assessed using the elevated plus maze (EPM) following our previously described protocol [36]. Each mouse was placed in the center of a plus-shaped maze elevated above the floor, consisting of two white open arms and two dark closed arms, and allowed to explore for 5 min. Sessions were recorded using a Basler GenICam camera and Media Recorder software (Noldus, VA, USA). The percentage of time spent in open and closed arms and the central zone, number of arm entries, total distance moved, and mean velocity were analyzed using EthoVision XT15 software (Noldus, VA, USA).

##### 2.5.3.4 Social interaction test

Social behavior was assessed using the three-chamber social interaction test as described previously by us [23,37,38]. The apparatus consisted of a Plexiglass box divided into three chambers with small openings between them. During habituation, the test mouse was allowed to explore all chambers freely for 10 min. Following habituation, the mouse was confined to the center chamber while a conspecific stranger mouse was placed inside a metal holding cage in one of the side chambers, and an identical empty cage was placed in the opposite chamber. The doors were then opened, allowing the test mouse to freely explore all chambers for 10 min. Behavior was recorded using a camera and Media Recorder software (Noldus, VA, USA). The total time spent interacting with the stranger mouse or empty cage, percentage of social interaction, and number of interaction bouts were scored by an observer blinded to the experimental groups. Heat maps representing social interaction patterns within each chamber were generated using EthoVision XT15 software. Representative heat maps are normalized for color within recordings, i.e. the same color is equally representative of relative spatial preference in all 3 images. Heat map generation was performed using the “base of tail” detection point, as this was found to be most representative of the animals’ detected position throughout the video recording.

### 2.6 *In-situ* hybridization (RNAscope)

#### 2.6.1 Visium DEG experiment

5 male mice of each genotype (C57, htau hemizygous) were anesthetized with avertin and fixed via transcardial perfusion of PBS followed by 4% PFA. Brains were cryoprotected in 30% sucrose before being embedded in OCT and flash frozen. Mouse brains were cryosectioned serially at 14 µm, sections were mounted on histobond glass slides, and RNAscope was performed according to the ACDbio manufacturer’s protocol using commercially-available probes for *Tph2* (cat. #318691) and *Slc17a8* (cat. #1199011) with either *Slc24a5* (cat. #1150641), *Kcna4* (cat. #405311), or *Scn4b* (cat. #484771). Images were acquired using an Olympus VS200 slidescanner at 20X. RNAscope analysis was performed using Qu-Path software (v 0.5) in accordance with the protocol outlined in Secci et al., 2023 [39]. A pixel classifier was first used to isolate *Tph2*+ area, and cellular detection was then performed for *Slc17a8* within this annotation, isolating cells that were double-positive for *Tph2* and *Slc17a8*. The subcellular detection function was then used to quantify transcript puncta for *Slc24a5*, *Kcna4*, or *Scn4b* in separate reactions. Validation of the subcellular detection algorithm was performed using manual puncta counts as suggested by the developer. Subcellular detection parameters were adjusted until the algorithm count was found to be within 5% of the manually-counted value, bounding a linear range from single cell to anatomical region. This validation was performed once for each genotype group for each gene, respectively.

#### 2.6.2 DREADDs fos experiment

2 htau mice from each of the DREADDs treatment groups (hM3Dq, hM4Di) and timepoints (acute CNO, 7-day CNO washout) were perfused, prepared, and assessed via RNAscope as in **Section 2.6.1** using commercially-available probes for *Tph2* (cat. #318691), *mCherry* (cat. #513201), and *Fos* (cat. #316921). Qu-Path software was used to perform the analysis. Cellular detection was performed for *Tph2* to identify 5-HT neurons, and object classifier were then used to categorize *Tph2*+ cells as either *Tph2*+, *Tph2*+ *mCherry*+ double-positive, *mCherry*+ *Fos*+ double-positive, *Tph2+ Fos*+ double-positive, or *Tph2+ mCherry+ Fos+* triple-positive. The percentage of *Tph2+ mCherry+* double-positive that were additionally *Fos+* triple-positive were calculated. 2 tissue sections were assessed per mouse, sampled from the extreme rostral/caudal regions of the injection location. Each data point in **Figure 4C** represents 1 tissue section, with each mouse contributing 2 separate data points.

**Figure 1.**
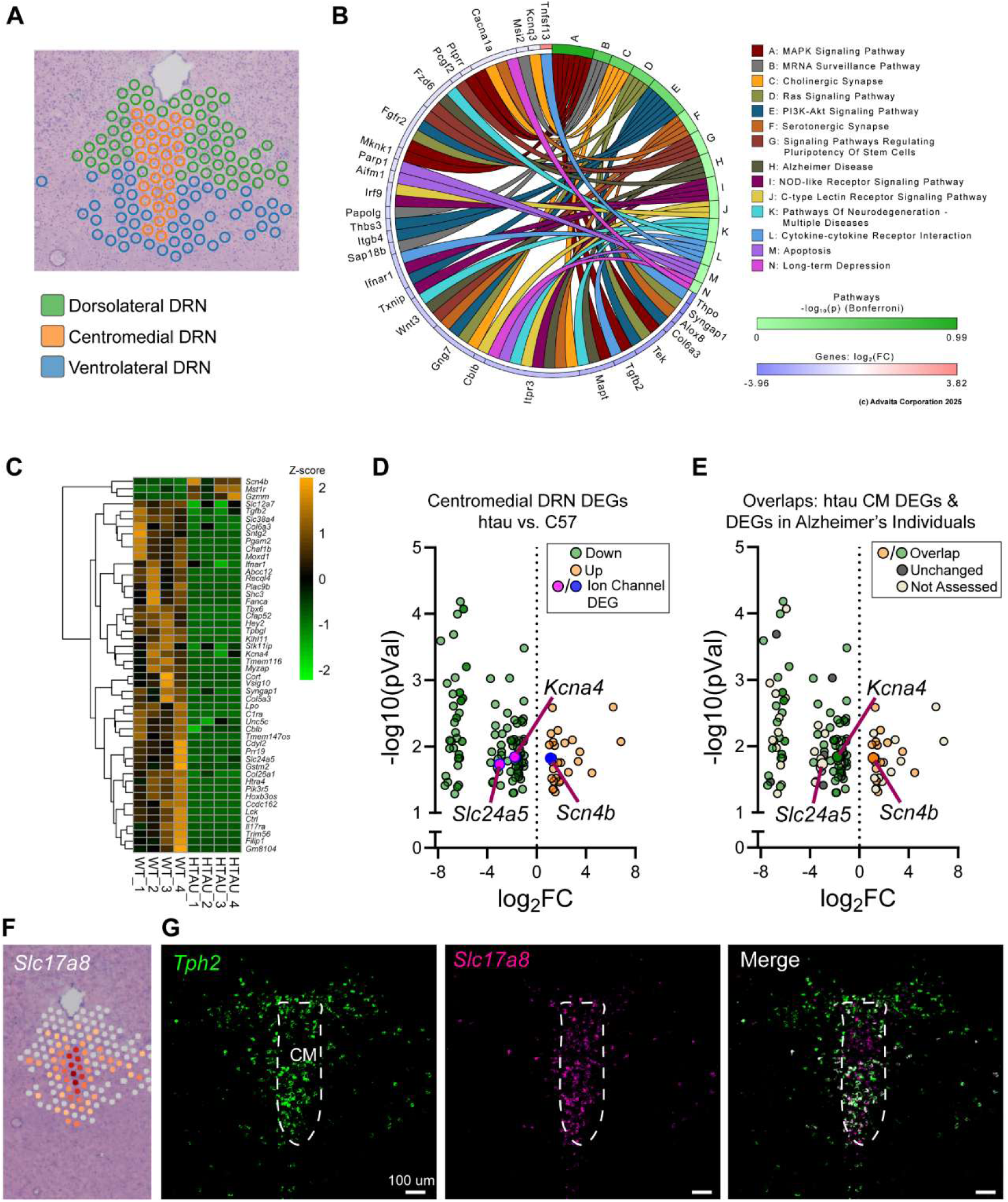
Differentially-expressed and marker genes of the centromedial dorsal raphe nucleus (DRN). ***A:*** Transcriptomically-distinct regions of the DRN as previously determined using Visium Spatial Gene Expression. ***B:*** Disease-relevant pathways affected by differentially-expressed genes (DEGs) in the centromedial DRN of htau mice as compared to wild-type mice. ***C:*** Hierarchical clustering of centromedial DEGs, top-50 delta only. Expression association is Z-scored for comparison across genes. ***D:*** Volcano plot of centromedial DRN DEGs, with ion channel genes annotated. ***E:*** Plot from (***D***), compared to the Agora database of human post-mortem Alzheimer’s DEGs. Genes that retain green/orange coloring were found to be differentially expressed in both Agora and our datasets, independent of their respective differential direction in human brain. ***F:*** Heatmap expression of *Slc17a8* (VGLUT3) within the DRN from our Visium dataset. ***G:*** RNAscope labeling of *Tph2*(green, left, marker for serotonin), *Slc17a8* (magenta, center), and both (right) in the centromedial (CM) DRN.

**Figure 2.**
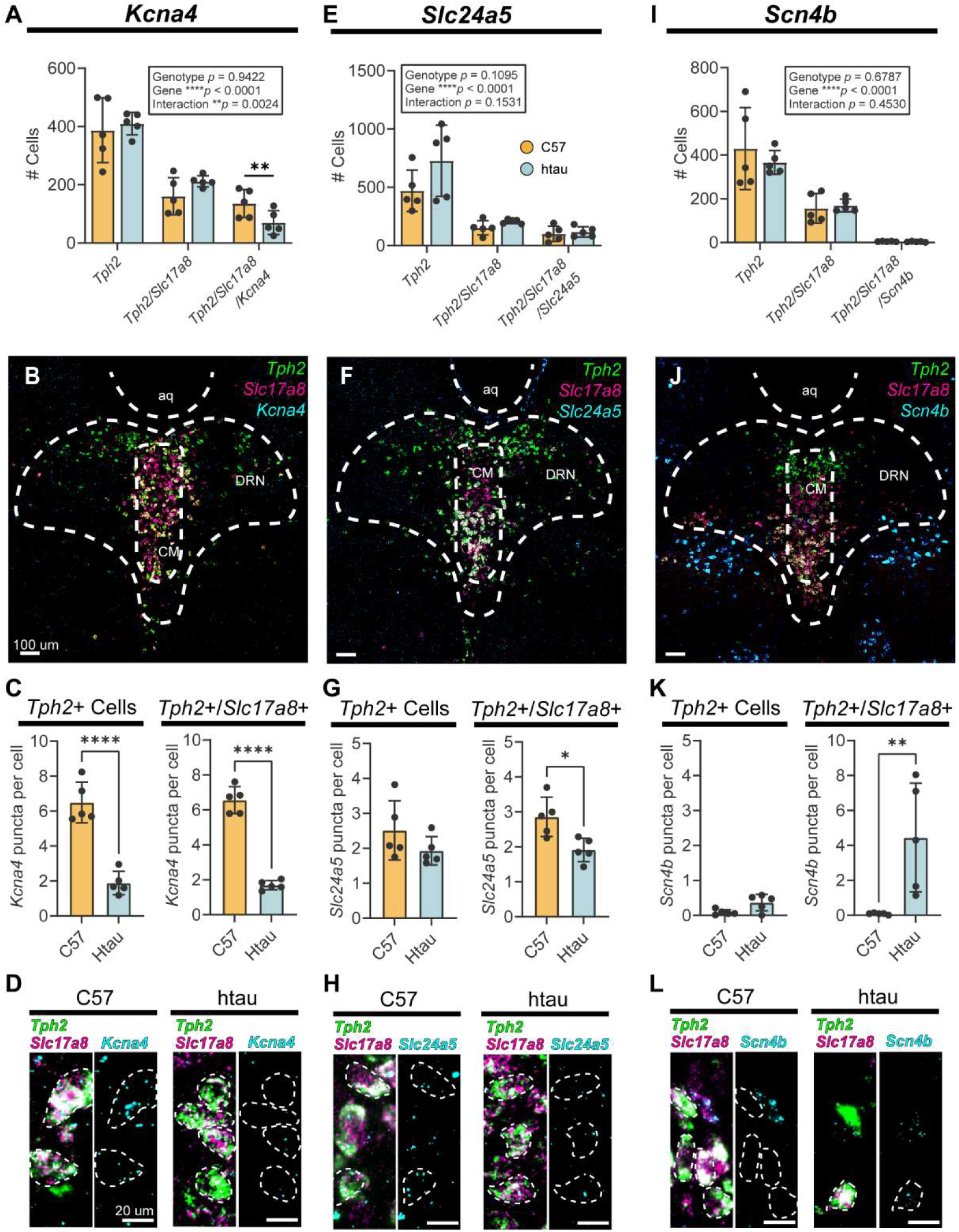
RNAscope validation of ion-channel DEGs in 5HT/glut neurons of the DRN. ***A:*** Number of serotonergic cells (*Tph2*), 5HT/glut cells (*Tph2*/*Slc17a8* (VGLUT3)) and *Kcna4*-(Kv1.4)-expressing 5HT/glut cells (*Tph2*/*Slc17a8*/*Kcna4*) in the DRN, wild-type (C57) and htau mouse, one tissue section per animal. The percentage of triple-positive cells, respective of double-positive cells, was found to be reduced in htau mice. ***B:*** Confocal image demonstrating the expression of *Tph2*, *Slc17a8*, and *Kcna4* with regards to the centromedial (<u>CM</u>) DRN; <u>aq</u>: aqueduct. ***C:*** Quantitation of *Kcna4* puncta within all serotonergic cells (*Tph2*+, left) and 5HT/glut cells (*Tph2*+/*Slc17a8*+, right) of the DRN. ***D:*** Representative images of *Kcna4* expression in 5HT/glut cells, C57 (left) and htau (right). ***E–H:*** same as ***A–D***, but for *Slc24a5* (NCKX5). ***I– L:*** same as ***A–D***, but for *Scn4b* (Navβ4). ***A,E,I:*** 2w RM ANOVA w/Sidak’s *post hoc*. ***C,G:*** Student’s *t*-test. ***K:*** Welch’s *t*-test (left), Mann-Whitney test (right). \**p* < 0.05, \*\**p* < 0.01, \*\*\*\**p* < 0.0001

**Figure 3.**
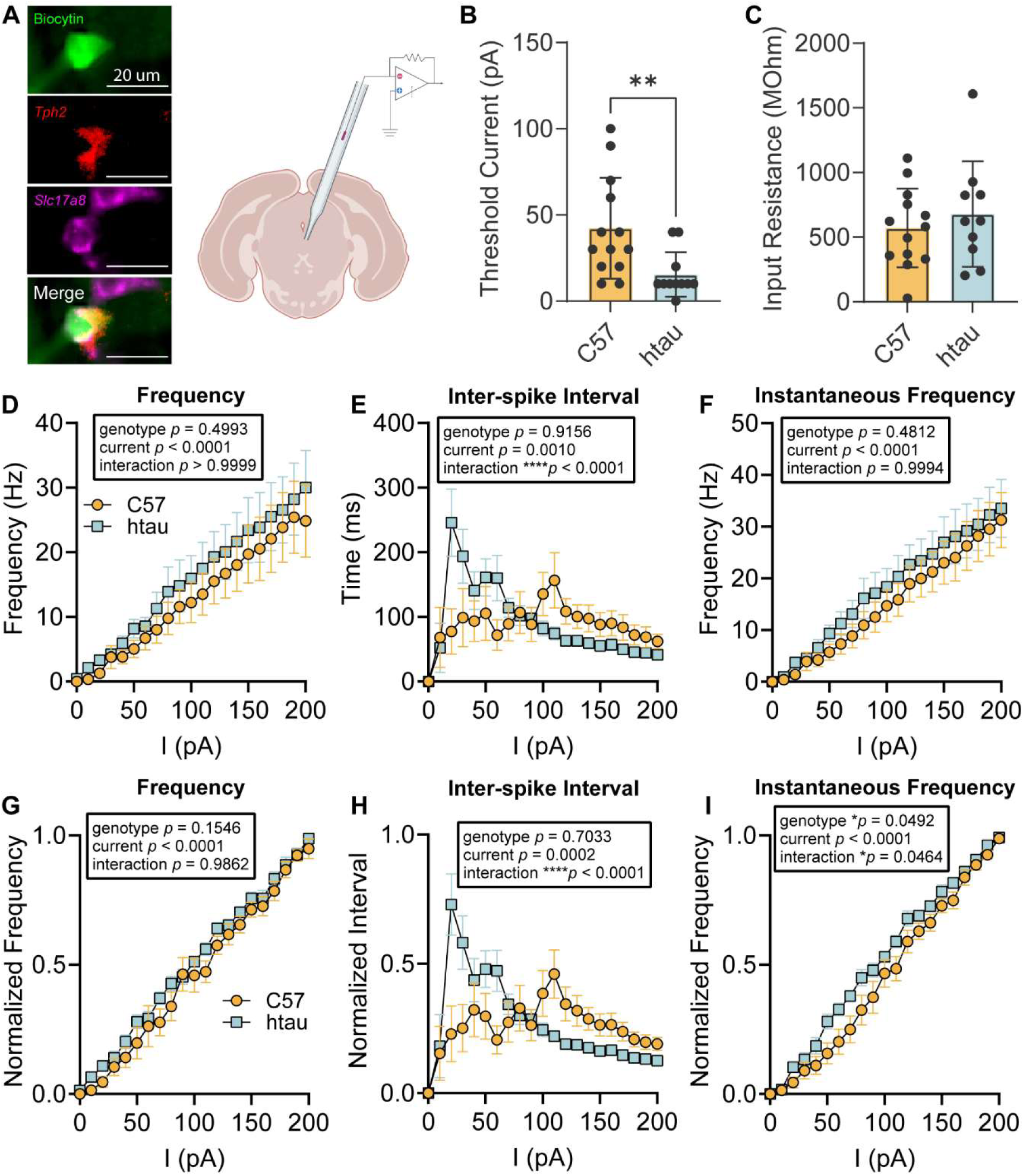
Changes in evoked excitability and action-potential df/dt of 5HT/glut neurons. ***A:*** *Post hoc* determination of 5HT/glut neurons using biocytin conjugation (green) in tandem with RNAscope targeting of *Tph2* (red) and *Slc17a8* (VGLUT3; magenta). ***B:*** Threshold current of 5HT/glut neurons, as determined using incremental current injections from -100–200 pA, P_d_ = 1 s, interval 10 s. ***C:*** Input resistance measurements obtained at -60 mV holding potential. ***D–F:*** Current-clamp recordings obtained using the recording paradigm in (***B***), C57 vs htau 5HT/glut neurons, demonstrating event frequency (***D***), average inter-spike interval (***E***), and instantaneous frequency (***F***). 2w mixed model with RM. ***G–I:*** Same as ***D–F***, but normalized within each recorded cell to compare current-dependent action-potential df/dt. ***B:*** Mann-Whitney test. ***C:*** Student’s *t*-test. ***D–I:*** 2w RM ANOVA w/Sidak’s *post hoc*. \**p* < 0.05, \*\**p* < 0.01, \*\*\*\**p* < 0.0001

**Figure 4.**
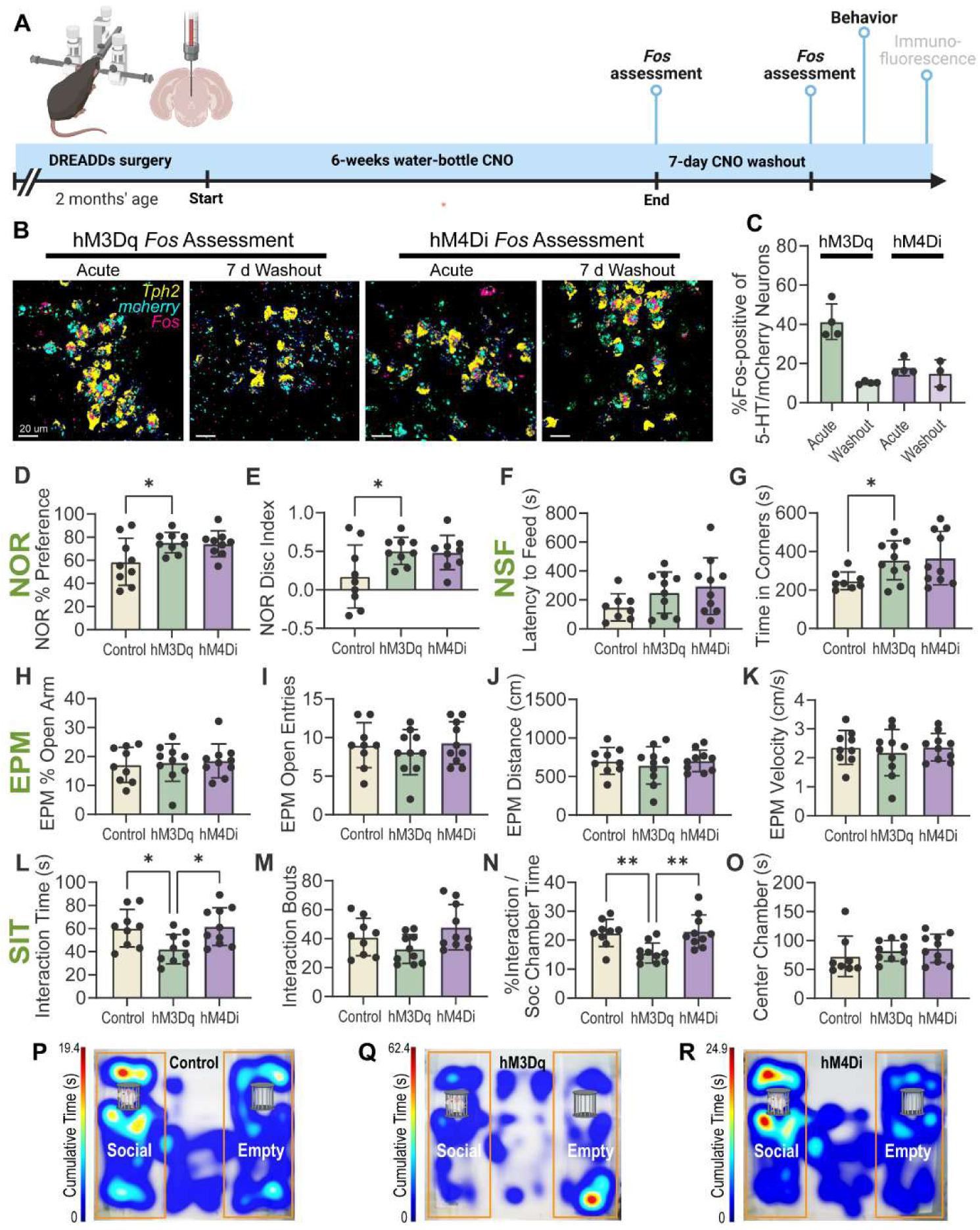
The effects of chronic CM DRN modulation on htau behaviors. ***A:*** Graphical overview of the htau DREADDs experiment. Htau mice were intracranially administered hM3Dq (n = 14), hM4Di (n=14), or control virus (n = 9) at 2 months’ age. DREADDs agonist (CNO DHC) was administered via water bottle for 6 weeks. ***B***,***C***: RNAscope assessment of *Fos* expression in virally transduced *Tph2*+ cells before CNO washout (acute) and after 7-day washout. Two data points were collected per mouse in (***C***). No statistical test was performed in (***C***), data were qualitatively compared. ***D,E***: Novel objection recognition (NOR). ***D***: Percent preference for novel object and ***E***, discrimination index. 1w ANOVA with Tukey’s *post hoc*. ***F,G***: Novelty-induced suppression of feeding (NSF). ***F***: latency to feed and ***G***, time spent in the corners of the arena during 10-minute assessment period. ***H–K***: Elevated plus maze (EPM) task. ***H***: Percent of total time spent in open arm, ***I***, entries to open arm, ***J***, total distance traveled, and ***K***, average velocity during the task. ***L–R***: Three-chamber social interaction test (SIT). ***L***: Total social interaction time, ***M***, number of social interactions, ***N***, social interaction time as a percentage of time spent in the social chamber, and ***O***, time spent in the center chamber. ***P–R*** Representative heat maps, normalized within trial. ***D–F,H–K***: 1w ANOVA w/ Dunnett’s *post hoc*. ***G***, 1w Welch’s ANOVA w/Dunnett’s *post hoc*. ***L-O***: 1w ANOVA w/ Tukey’s *post hoc*.\**p* < 0.05, \*\**p* < 0.01

### 2.7 Immunofluorescence (IF)

After the conclusion of behavioral assessment, all remaining mice from the DREADDs experiment (9 control, 10 hM3Dq, 10 hM4Di) were perfused and brains were harvested as in **Section 2.6.1**. Brains were cryosectioned serially at 20 µm and stored in anti-freeze medium until IF. Coronal sections containing the DRN were mounted on histobond glass slides, blocked for 60’, and incubated in primary antibodies targeting Tph2 (goat, Everest EB11012, 1:375), VGLUT3 (mouse, Sigma-Aldrich SAB5200312, 1:500) phosphorylated human tau (pTau; AH36, rabbit, StressMarq SMC-601, 1:500) at 4°C overnight. Species-specific secondary antibodies were incubated for 2 hours at room temperature. Blocking buffer consisted of 10% NDS, 0.5% Triton X-100, and 0.2% Tween-20 in PBS. Incubation buffer consisted of 2% NDS and 0.1% Tween-20 in PBS. Confocal z-stacks (1 µm) were captured using an Olympus FV3000 laser scanning confocal microscope (6 sections/z-stack) and converted to maximum-intensity projection images using FIJI (ImageJ) software. Percent-immunoreactive area (% IR) calculations were then performed using Qu-Path software. The DRN and CM DRN were manually bounded using the aqueduct and Tph2 distribution as landmarks. A pixel classifier was then created for the mCherry image channel within each ROI. The percentage of each ROI that was measured to be positive for mCherry was used to determine “virus coverage” represented in **Figure 5H,I**. An annotation was created for the mCherry pixel classifier, and a second pixel classifier was nested within the mCherry-positive area using the AH36 (pTau) image channel. The percentage of mCherry-positive area that was found to be dually pTau-positive is represented as “pathology” in **Figure 5H,I**.

**Figure 5.**
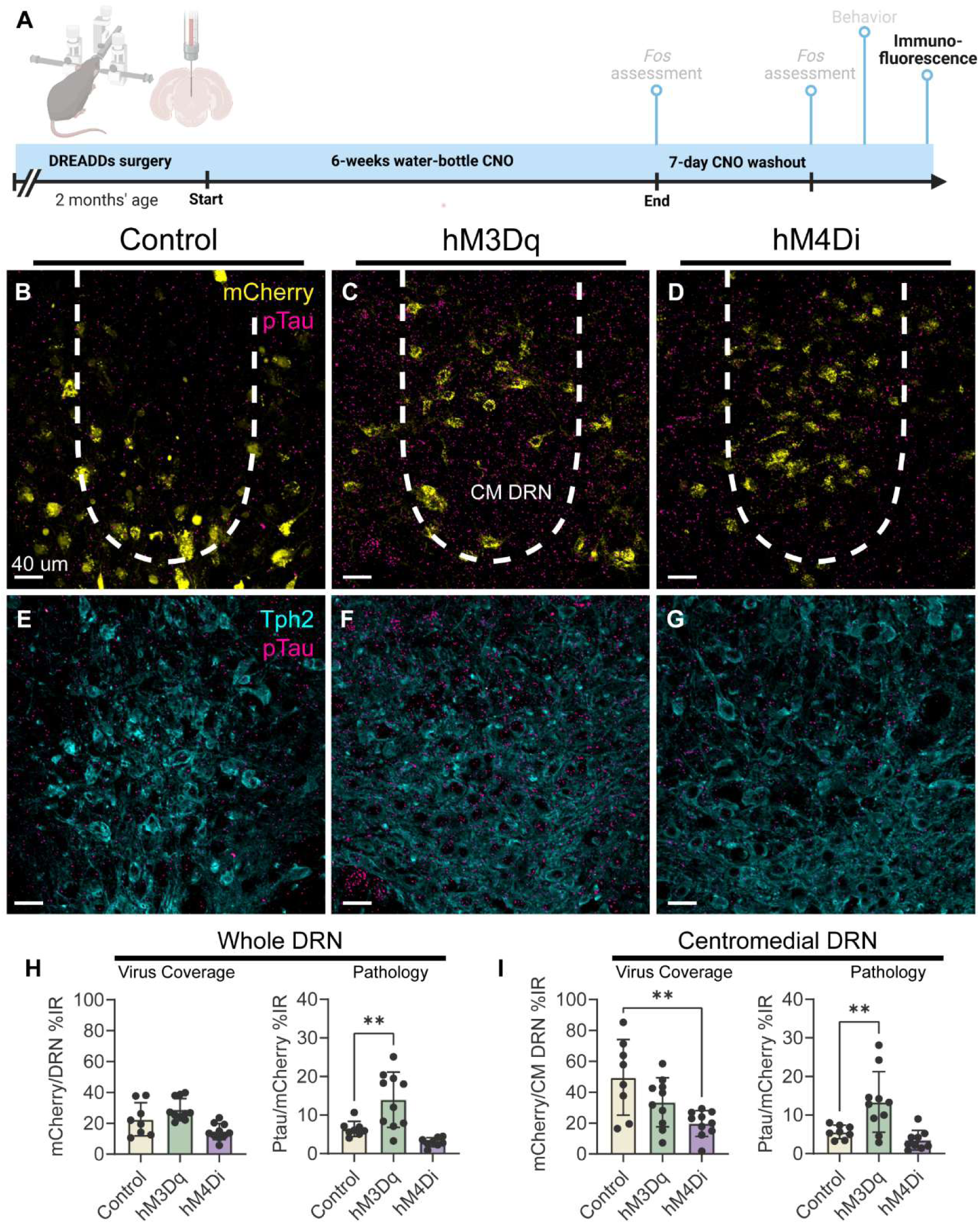
Chronic hyper-excitability enhances accumulation of pTau within the DRN of htau mice. ***A:*** Graphical overview of the htau DREADDs experiment. ***B–G:*** Confocal images of the DRN with immunolabeled Tph2 and pTau (AH36 antibody). Viral mCherry reporter (***top row***) was not amplified before imaging. One section per mouse that contained virally transduced cells was stained and imaged. ***H***, ***left:*** Comparison of total virus coverage within the DRN expressed as percent immunoreactive area (<u>%IR</u>). Measurements were made for all mCherry+ area, Tph2 staining was used for anatomical alignment only. ***H, right:*** Assessment of pathology within the DRN, expressed as % IR for pTau immunolabeling within mCherry+ pixels of the DRN. ***I:*** Same as (***H***), but limited to the CM DRN. 1w ANOVA with Dunnett’s *post hoc*. \*\**p* < 0.01

### 2.8 Electrophysiology

#### 2.8.1 Brain slice preparation

This procedure has been described in detail previously [23]. Briefly, deeply anesthetized mice were transcardially perfused with ice-cold oxygenated modified artificial cerebrospinal fluid. Brains were quickly dissected, and coronal slices containing the dorsal raphe nucleus (DRN, 300 μm) were obtained using a vibratome (VT1200S; Leica Biosystems, Wetzlar, Germany).

#### 2.8.2 *Ex vivo* electrophysiological recordings

Neurons were visualized using an upright microscope (BX51W1; Olympus, Tokyo, Japan) accompanied by a differential interference contrast imaging system. Membrane currents were amplified with a Multiclamp 700 B amplifier (Molecular Devices, San Jose, CA, USA), filtered at 3 kHz, and sampled at 20 kHz with a Digidata 1550B digitizer (Molecular Devices). Data were acquired via the pClamp 11 software (Molecular Devices). Access resistance was monitored and changes greater than 20% would lead to discontinuation of the recordings. After establishing a whole-cell configuration, cells were first sampled for proper access resistance (less than 40 MΩ) prior to initiating a series of current-clamp recordings. Cells were held at -55 mV between pulses to compare data across cells. Peri-threshold current levels were determined by incrementally injecting a family of current pulses of 1000 ms duration. Current was injected every 10 s and was stepped in 10 pA increments from -100 pA to 200 pA to obtain action potential characteristics and current-dependence data. Input resistance was calculated using the voltage deflection elicited by a current injection of -10 pA.

#### 2.8.3 Verification of cellular identify

To limit data analyses to dual serotonergic/glutamatergic neurons, *post hoc* confirmation of cellular identity was performed using intracellular recording solution containing biocytin. After obtaining recordings, slices were fixed in 4% PFA overnight. Slices were then embedded in OCT, cryosectioned at 20 µm, and mounted on histobond glass slides. The slides were incubated with streptavidin conjugated to Alexa 488 overnight, then RNAscope was performed targeting *Tph2* and *Slc17a8*. The proteinase digestion step of the RNAscope protocol was omitted to preserve biocytin/streptavidin labeling. Images were acquired using an Olympus VS200 slidescanner at 20X, and co-labeling was visually assessed using Qu-Path software (0.5).

### 2.9 Post-mortem human AD tissue procurement

Formalin-fixed, paraffin-embedded (FFPE) tissue samples were obtained from the Iowa Neuropathology Resource Laboratory (INRL) at the University of Iowa. Details of individual cases, including demographic information and sources, are provided in Table 1. All samples were reviewed by an experienced neuropathologist (Dr. Marco Hefti) to confirm the presence of appropriate sections of the dorsal raphe nucleus and tau pathology. The University of Iowa Institutional Review Board determined that this study does not constitute human subjects research under the NIH Common Rule, as it exclusively used tissue from deceased individuals (determination #201706772). All procedures were performed in accordance with institutional, national, and international ethical standards, including the 1964 Declaration of Helsinki and its subsequent amendments or equivalent guidelines.

**Table 1.** Clinical and demographic data for human cases. Tissue samples were procured from the Iowa Neuropathology Resource Laboratory (INRL). Ages not found to differ between groups (CN 61.3 ± 7.9 yrs vs AD 65.4 ± 3.4 yrs, *p* = 0.3220). All tau-positive cases were included in the analyses of Figure 6, all cases were included in the analyses of Figure 7.

| INRL Case ID | Age | Sex | DRN Tau Pathology | Diagnosis | Braak Stage | Cognitive Status | Data in Figure |
| --- | --- | --- | --- | --- | --- | --- | --- |
| A23-176 | 53 | M | - | Control | - | CN | 6(G,H), 7 |
| A24-056 | 56 | M | - | PART | - | CN | 6(G,H), 7 |
| A23-288 | 67 | M | + | Control | - | CN | 6, 7 |
| A20-332 | 69 | M | + | Control | - | CN | 6, 7 |
| A24-250 | 62 | M | + | AD/LBD | B2 | AD | 6, 7 |
| A24-297 | 62 | M | + | AD | B2 | AD | 6, 7 |
| A23-119 | 66 | M | + | AD | B2 | AD | 6, 7 |
| A23-282 | 67 | M | + | AD/LBD | B2 | AD | 6, 7 |
| A23-319 | 70 | M | + | AD/LBD | B2 | AD | 6, 7 |

### 2.10 In Situ Hybridization (FISH) for Post-Mortem Human AD Tissue

Formalin-fixed, paraffin-embedded (FFPE) tissue blocks were sectioned at 10 µm and mounted directly onto Slidemate Laser slides. Multiplexed fluorescence in situ hybridization (FISH) was performed using a modified ACDbio RNAscope® V2 protocol with Pretreat-Pro reagent (Cat. No. 323100). Briefly, slides were baked for 1 hour at 60 °C, deparaffinized, dehydrated, and treated with hydrogen peroxide. Target retrieval was performed for 25 minutes using the manufacturer’s reagent, followed by incubation with Pretreat-Pro reagent for 35 minutes at 40 °C in a HyBez™ Hybridization Oven. After pretreatment, mRNA probes targeting *TPH2* (#C1-25129a), *SLC17A8* (#C2-25129b), *KCNA4* (#C3-888751), or *SLC24A5* (#C3-1674181) were applied. Subsequent amplification and Opal detection steps were carried out according to the manufacturer’s instructions. Sections were treated with TrueVIEW® Autofluorescence Quenching reagent (Vector Laboratories, CA, USA) for 3 minutes, then mounted with ProLong™ Gold Antifade Mountant (P36930, Thermo Fisher Scientific). Slides were imaged using an Olympus SLIDEVIEW VS200 digital slide scanner, and images were analyzed in QuPath (version 0.5.1). A pixel classifier was trained to detect the *TPH2*⁺ area, within which *SLC17A8*⁺ objects were identified. The intensity of *SLC17A8* expression within *TPH2*⁺ neurons was quantified. Within the dual *TPH2*⁺/*SLC17A8*⁺ annotation, *KCNA4*⁺ or *SLC24A5*⁺ objects were identified, and their expression intensities were measured. The percentage of *KCNA4* and *SLC24A5* mRNA expression area (EA) was calculated within both *TPH2*⁺/*SLC17A8*⁺ and non-*TPH2*⁺/*SLC17A8*⁺ neurons.

### 2.11 Immunofluorescence-Enabled FISH (iFISH) for Post-Mortem Human AD Tissue

To enable detection of pathologically phosphorylated tau (pTau) within serotonergic/glutamatergic neurons in post-mortem AD and control tissue, fluorescence in situ hybridization was combined with immunofluorescence (iFISH). FISH was performed as described above (Section 2.10), followed by immunofluorescence prior to the quenching step. FISH-processed slides were washed briefly in 0.01 M PBS, permeabilized with 0.5% Triton X-100/PBS for 30 minutes, and blocked in 10% normal donkey serum (NDS) with 0.1% Triton X-100 in PBS. Slides were then incubated overnight at 4 °C with primary antibodies against MAP2 (chicken, #NB300-213, Novus Biologicals; 1:200) and phosphorylated tau (AT8, mouse, #MN1020, Thermo Fisher Scientific; 1:100). The following day, species-specific Alexa Fluor– conjugated secondary antibodies (Jackson ImmunoResearch Laboratories) were applied for 2 hours at room temperature. After secondary incubation, sections were treated with TrueVIEW® reagent to quench tissue autofluorescence and mounted with ProLong™ Gold Antifade Mountant. Imaging was performed using an Olympus FV3000 confocal microscope with z-stacks (0.5 µm/step) at 20× or 40× magnification (optical resolution 1× or 2×). Multiple tiled images were captured per region of interest, and mosaics were generated using FV3000 Multi-Area Time-Lapse (MATL) software. Confocal stacks were converted to maximum projection images in QuPath (version 0.5.1). Using a pixel classifier, *TPH2*⁺ regions were identified and annotated. Within these, *SLC17A8*⁺ objects were segmented, followed by identification of AT8⁺ objects within dual *TPH2*⁺/*SLC17A8*⁺ regions. The intensity of AT8 immunoreactivity within *TPH2*⁺, *TPH2*⁺/*SLC17A8*⁺, and non-*TPH2*⁺/*SLC17A8*⁺ neurons was quantified, and the percentage of AT8 immunoreactive area (IR) was calculated for each population.

### 2.12 Experimental design and statistical analyses

Data analyses and statistical tests were performed using Prism version 10 (GraphPad Software, Inc., La Jolla, CA, USA; RRID:SCR_002798). Visium Spatial Gene Expression data were analyzed as described previously [25]. Sample sizes for mouse RNAscope reactions were estimated using the log fold-change and variation from the Visium dataset. Two-way comparisons (electrophysiology data, **Figure 3D-I**) use a 2-way mixed model with repeated measures (2w ANOVA). Normalized plots in **Figure 3G-I** are normalized to the peak value within each respective recording. Pairwise comparisons and one-way ANOVA for all other experiments were performed as follows: outliers were first determined using a ROUT test (Q = 1%). Data were then checked for normal distribution using a Shapiro-Wilk test, and homogeneity of variance using the Fmax test. Where data violated homogeneity of variance, a non-parametric analysis was used. All DREADDs experiments (**Figure 4,5**) use 1w ANOVA. Bar charts annotate means, with plungers representing standard deviation. 2-way plots represent means with plungers representing standard error.

## 3 RESULTS

### 3.1 The mouse centromedial DRN is characterized by serotonin neurons that are dually glutamatergic

Using the list of differentially-expressed genes (DEGs) that we previously identified [25] in the centromedial DRN of htau mice (**Figure 1A**), Gene Ontology analysis identified several affected pathways relevant for MAPK signaling, Alzheimer’s disease, serotonergic synapse, long-term depression, and synaptic function (**Figure 1B**). Many DEGs were implicated across multiple pathways, particularly three ion channel-related genes: *Kcna4* (Kv1.4; potassium voltage-gated channel subfamily A member 4), *Slc24a5* (NCKX5; sodium/potassium/calcium exchanger 5), and *Scn4b* (Navβ4; sodium voltage-gated channel beta subunit 4) (**Figure 1C,D**). We next compared this list of centromedial DRN DEGs to the Agora gene expression database, which catalogs RNA expression findings of human Alzheimer’s experiments involving the anterior cingulate cortex, cerebellum, dorsolateral prefrontal cortex, frontal pole, inferior frontal gyrus, posterior cingulate cortex, parahippocampal gyrus, superior temporal gyrus, and temporal cortex. Though Alzheimer’s-related changes in gene expression are not catalogued for the dorsal raphe nucleus, 46 of our centromedial DEGs were found to be Agora DEGs, 7 of our DEGs were not Agora DEGs, and the remaining 43 DEGs of our centromedial dataset were not previously assessed in AD (**Figure 1E**). Of primary importance, *KCNA4* and *SCN4B* were found to be differentially expressed in multiple Alzheimer’s brain regions, with *SLC24A5* not yet assessed.

Given that the anatomical organization of the dorsal raphe nucleus differs between the murine and human brain, we sought to identify “marker genes” that would enable more precise identification of 5-HT neurons within the centromedial DRN. We used the Loupe Browser’s differential expression tool to identify genes that are differentially expressed within the centromedial DRN as compared to other DRN subregions. Interestingly, we found that *Slc17a8* (VGLUT3, a marker for glutamatergic cells) was highly enriched in the centromedial DRN (**Figure 1F**). Using RNAscope, we confirmed the presence of dual serotonergic glutamatergic (5HT/glut) neurons that were found to be primarily confined to the centromedial DRN (**Figure 1G**). Within the mouse DRN, we found that ∼40% of 5-HT neurons are dually glutamatergic, with no apparent difference between animal genotypes (C57 39.4 ± 9.4 % vs htau 43.9 ± 11.4 %, *p* = 0.1096).

### 3.2 Htau 5HT/glut neurons differentially express ion-channel genes

We next used RNAscope to validate the differential expression of these ion channel-related genes in 5HT/glut neurons. These comparisons were performed in male C57 mice vs htau mice aged 16 weeks, as we performed our original Visium Spatial Gene Expression reaction using these experimental conditions [25]. We used RNAscope probes for *Tph2* (marker for serotonin production) and *Slc17a8* (VGLUT3) to identify 5HT/glut neurons, and each DEG was investigated in separate reactions, respectively. Within the DRN, we found *Slc17a8* to be expressed in ∼41% of *Tph2*-positive neurons, with no difference C57 vs htau (*p* = 0.1096). We next examined expression of *Kcna4*, which was expressed in a greater percentage of C57 5HT/glut neurons as compared to htau 5HT/glut neurons (**Figure 2A,B**, C57 85.50 ± 7.52 % vs htau 32.94 ± 19.47%, \*\**p* = 0.0079). Expression of *Kcna4* was found to be reduced in DRN 5-HT neurons overall (**Figure 2C left**, C57 6.495 ± 1.167 puncta/cell vs htau 1.889 ± 0.665 puncta/cell, \*\*\*\**p* < 0.0001) and also in DRN 5HT/glut neurons (**Figure 2C right**, C57 6.559 ± 0.767 puncta/cell vs htau 1.694 ± 0.265 puncta/cell, \*\*\*\**p* < 0.0001; **Figure 2D**). Within the DRN, we found *Slc24a5* to be expressed in ∼59% of 5HT/glut neurons, with no difference C57 vs htau (**Figure 2E,F**). Expression of *Slc24a5* was not found to be reduced in DRN 5-HT neurons overall (**Figure 2G left**), but was found to be reduced in DRN 5HT/glut neurons (**Figure 2G right**, C57 2.859 ± 0.562 puncta/cell vs htau 1.913 ± 0.336 puncta/cell, \**p* = 0.0120; **Figure 2H**). We found *Scn4b* to be expressed in only ∼3% of DRN 5HT/glut neurons, with no difference C57 vs htau (**Figure 2I,J**). Expression of *Scn4b* was not found to be increased in DRN 5-HT neurons (**Figure 2K left**), but was increased in DRN 5HT/glut neurons, albeit with high variability (**Figure 2K right**, C57 0.089 ± 0.047 puncta/cell vs htau 4.450 ± 3.112 puncta/cell, \**p* = 0.0351; **Figure 2L**).

### 3.3 Htau 5HT/glut neurons have altered biophysical properties and firing patterns

Given the known roles of Kv1 channels (shaker potassium channels) in neuronal excitability and the refractory phase of action potentials, we next used *ex vivo* electrophysiology to assess these properties in male C57 and htau mice aged 16 weeks. Using biocytin-spiked intracellular recording solution in tandem with RNAscope, we verified the identity of recorded neurons *post hoc* (**Figure 3A**). We observed that 5HT/glut neurons of htau mice are more excitable, requiring smaller stimuli to evoke action potentials (rheobase) (**Figure 3B**, C57 42.31 ± 29.20 pA vs htau 15.45 ± 12.93 pA, \*\**p* = 0.0055). We observed no difference in input resistance, C57 vs htau (**Figure 3C**). We next examined action potential frequency and df/dt (rate of change of frequency) by assessing event frequency, inter-spike interval, and instantaneous frequency (the fastest occurrence between spikes) during 1 s current injections, stepped from -100 to +200 pA in increments of 10 pA (**Figure 3D–F**). Overall, we found that htau 5HT/glut neurons tend to fire more intermittently than C57 in response to smaller stimuli (less than 90 pA), and more continuously in response to larger stimuli (**Figure 3E,H**, \*\*\*\**p* < 0.0001), with no difference in event frequency (**Figure 3D,G**). We further found that htau 5HT/glut neurons exhibited faster normalized instantaneous frequency vs C57 (**Figure 3I**), both overall (\**p* = 0.0492) and across changes in current magnitude (\**p* = 0.0464).

### 3.4 Chronic hyper-excitability affects 5-HT-related behaviors and enhances pTau accumulation within the DRN of htau mice

The htau mouse is commonly used to model the prodromal phase of AD [40], due to its slow rate of pathology development [41]. Because we found enhanced excitability and action-potential firing in htau 5HT/glut neurons, we sought to determine whether magnifying this phenotype would affect 5-HT-related behaviors or the accumulation of pathologically-phosphorylated tau (pTau). To accomplish this, we used *in vivo* chemogenetics tools (Designer Receptors Exclusively Activated by Designer Drugs; DREADDs). At 8 weeks’ age, male htau mice were stereotaxically injected with AAV targeting the centromedial DRN, carrying either an excitatory (AAV8-hM3Dq-mCherry), inhibitory (AAV8-hM4Di-mCherry) or control (AAV8-mCherry) construct. After surgical recovery, mice were then administered the DREADD ligand clozapine-N-oxide dihydrochloride (CNO DHC) via water bottle in their home cage for 6 weeks. At 16 weeks’ age, mice underwent a 7-day “CNO washout” to abolish the DREADD effect on neuronal activity, isolating any changes in pTau accumulation as primary drivers for behavioral changes. **Figure 4A** contains a graphical overview of this experimental timeline. Two mice per DREADDs group were sacrificed at the beginning and end of the washout period to assess *Fos* expression in virally transduced cells (**Figure 4B,C**). After the 7-day CNO washout, mice underwent behavioral testing for DRN-related behaviors using the novel object recognition (NOR), novelty-induced suppression of feeding (NSF), elevated plus maze (EPM) and social interaction (SI) tests. In the chronic excitatory group, we observed an increased percent-preference for a novel object (**Figure 4D**, Control 58.69 ± 20.36 % vs hM3Dq 75.31 ± 8.85 %, \**p* = 0.0396) and enhanced discrimination index (**Figure 4E**, Control 0.174 ± 0.407 vs hM3Dq 0.506 ± 0.177, \**p* = 0.0396). During the NSF test, we observed no change in latency to feed (**Figure 4F**), but the chronic excitatory group spent more time in the corners of the arena (**Figure 4G**, Control 247.9 ± 45.4 s vs hM3Dq 354.3 ± 101.5 s, \**p* = 0.0313). We observed no differences between groups for any metrics of the EPM test (**Figure H–K**). During the three-chamber SI test, the chronic excitatory group exhibited a reduction in social interaction time (**Figure 4L**, Control 60.22 ± 16.28 s vs hM3Dq 42.10 ± 12.63 s, \**p* = 0.0387) with no difference in the number of social interaction bouts (**Figure 4M**). The percent of total time spent in the social chamber that was spent interacting with the stranger mouse was also decreased in the chronic excitatory group (**Figure 4N**, Control 22.52 ± 4.68 % vs hM3Dq 15.54 ± 3.45 %, \*\**p* = 0.0089); no difference was observed for time spent in the center chamber (**Figure 4O**).

Immediately following the behavioral assessments, we used immunolabeling to assess any changes in pTau accumulation resultant to the chronic DREADDs challenge (**Figure 5A**). Using antibodies targeting Tph2 and pTau (AH36 antibody) (**Figure 5B–G**), we determined that chronic hyperexcitability via hM3Dq enhanced the accumulation of pTau as compared to the Control group, measured via percent immunoreactive area (%IR) (**Figure 5H right,** Control 6.412 ± 1.973 %IR vs hM3Dq 13.99 ± 7.187 %IR, \*\**p* = 0.0044) with no apparent difference in overall virus coverage (**Figure 5H left**). In the CM DRN, we again found that hM3Dq enhanced the accumulation of pTau (**Figure 5I right**, Control 5.602 ± 1.737 %IR vs hM3Dq 13.380 ± 7.848 %IR, \*\**p* = 0.0070), with only a reduction in CM DRN virus coverage in the hM4Di group (**Figure 5I left**, Control 49.57 ± 24.48 %IR vs hM4Di 19.69 ± 8.37 %IR, \*\**p* = 0.0027). We found no difference in pTau/mCherry %IR with respect to DRN vs CM DRN (2w ANOVA *p* = 0.8259), and no difference in the percentage of VGLUT3+ neurons that were dually mCherry+ with respect to Control vs hM3Dq (Mann-Whitney *t*-test, Control 16.65 ± 10.29% vs hM3Dq 19.66 ± 9.48%, *p* = 0.7302).

### 3.5 5HT/glut neurons are especially vulnerable to tau pathology in human AD, and preferentially express *KCNA4* and *SLC24A5*

We next sought to determine whether our findings in htau 5HT/glut neurons mirrored those in post-mortem clinical AD. With assistance from the Iowa Neuropathology Resource Laboratory, we selected tissue sections containing the DRN from cognitively-normal (CN) controls and patients with a previous clinical diagnosis of AD. Tissue samples from CN controls were not excluded based on the presence of tau pathology, as we have previously found that ∼40% of CN control tissue contains tau pathology in the DRN [42]. Tissue samples from AD patients were not excluded based upon the co-diagnosis of Lewy body dementia, given that ∼50% of AD patients are co-diagnosed [43–45]. All AD cases were matched for Braak stage 2 in order to minimize variation in neurodegenerative progression. Ages were not found to differ between groups (CN 61.3 ± 7.9 yrs vs AD 65.4 ± 3.4 yrs, *p* = 0.3220).

Of four non-AD cases, two were found to contain tau pathology in the DRN. Thus, we first assessed all cases, independent of cognitive status, to determine any pathological or gene-expression differences in 5HT/glut neurons vs 5HT-nonglut neurons (5-HT neurons negative for *SLC17A8*). Each tissue sample contributed one data point each from its 5-HT, 5HT/glut and 5HT-nonglut neurons, respectively. We sought to determine the spatial localization of 5HT/glut neurons in the human DRN, and found that they are not confined to a discrete anatomical subregion. Instead, they are positioned throughout the DRN (**Figure 6A–C**). Using Immunofluorescence-enabled *in situ* hybridization (iFISH), we immunolabeled MAP2 to visualize neuronal boundaries and AT8 antibody to label pathological tau, in tandem with RNAscope probes targeting *TPH2* and *SLC17A8* (**Figure 6D**). We found 5HT/glut neurons to be especially vulnerable to tau pathology as compared to 5HT-nonglut neurons, as measured via AT8 %IR (**Figure 6F**, 5HT/glut 73.71 ± 7.42 % vs 5HT-nonglut 48.95 ± 10.54 %, \*\**p* = 0.0043). The expression of *KCNA4* and *SLC24A5* within the human DRN was confirmed by a previously-reported Microarray study in the ALLEN Brain Atlas Data Portal (**Figure 6E**) [46]. Using RNAscope, we further found both *KCNA4* and *SLC24A5* to be preferentially expressed in 5HT/glut neurons (**Figure 6G**, *KCNA4* 5HT/glut 52.27 ± 11.41 % vs 5HT-nonglut 7.37 ± 4.42 %, \*\*\*\**p* < 0.0001; **Figure 6H**, *SLC24A5* 5HT/glut 43.69 ± 14.28 % vs 5HT-nonglut 2.54 ± 2.99 %, \*\*\*\**p* < 0.0001).

**Figure 6.**
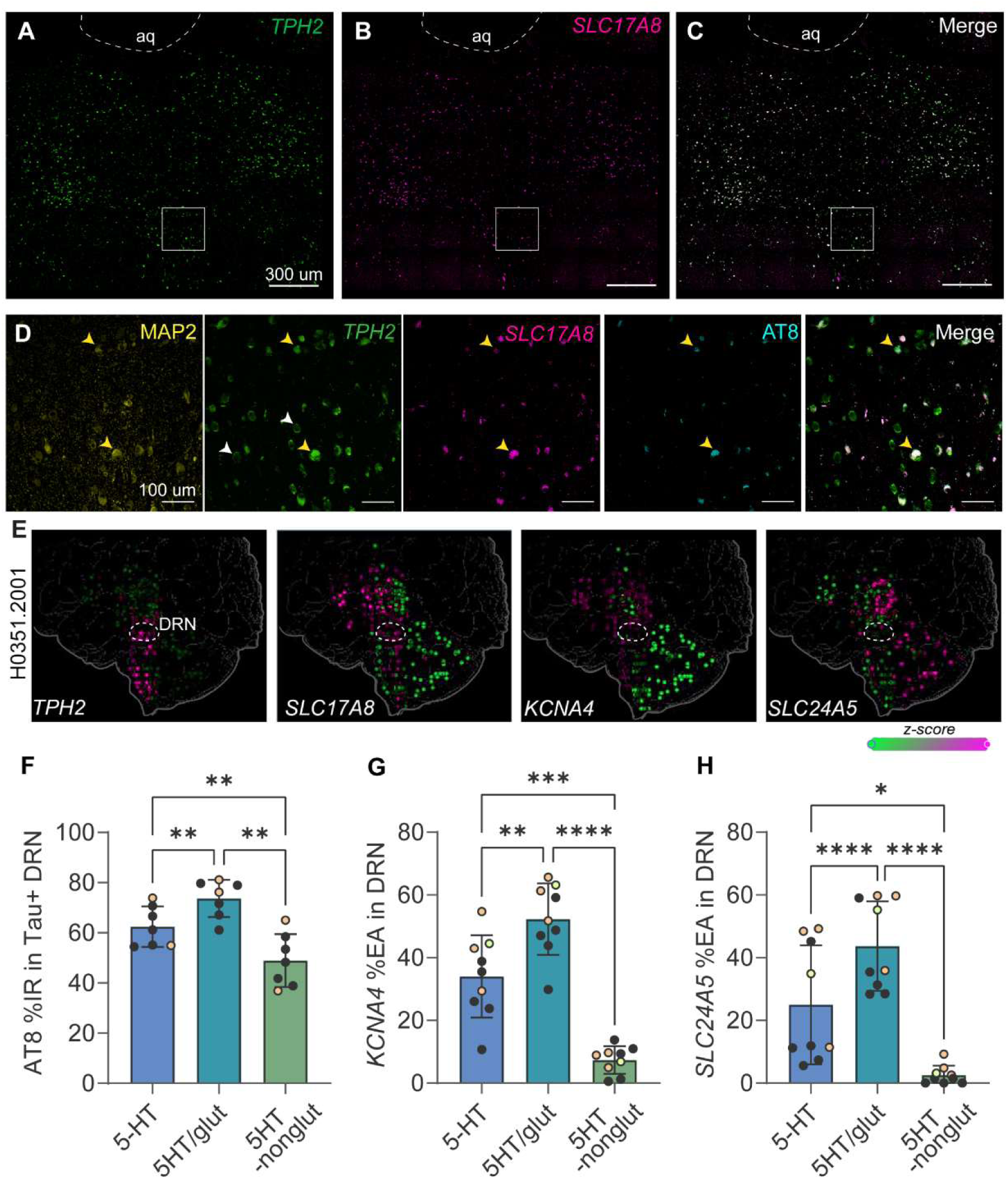
Anatomical, pathological, and gene-expression characteristics of human 5HT/glut neurons. ***A–C:*** Confocal images demonstrating spatial distribution via RNAscope labeling for *TPH2* (***A***), *SLC17A8* (VGLUT3) (***B***) and 5HT/glut neurons (***C***) in human DRN; <u>aq</u>: aqueduct. Annotated box is expanded in (***D***). ***D:*** Immunofluorescence-enabled *in situ* hybridization (iFISH) labeling for MAP2 (microtubule-associated protein 2), *TPH2*, *SLC17A8*, and AT8 (phospho-tau antibody). <u>Yellow arrows</u>: 5HT/glut; <u>white arrows</u>: 5HT-nonglut. ***E:*** Micro-array data for subcortical brain areas from the ALLEN Human Brain Atlas for *TPH2*, *SLC17A8*, *KCNA4*, and *SLC24A5* (Case #H0351.2001). ***F:*** Percent immunoreactive area (<u>%IR</u>) for AT8 within all *TPH2*-positive area (5-HT), *TPH2*-positive / *SLC17A8*-positive area (5HT/glut) and *TPH2*-positive / *SLC17A8*-negative (5HT-nonglut) area for all tau-positive human cases. <u>Orange data points</u> denote cognitively-normal clinical status. ***G:*** Percent expression area (<u>%EA</u>) for *KCNA4* within 5-HT, 5HT/glut and 5HT-nonglut areas, respectively. Dataset includes all cases; <u>lime data point</u> denotes cognitively-normal PART case (tau-negative DRN). ***H:*** Same as (***G***), but for *SLC24A5*. ***F–H:*** 1w RM ANOVA w/Tukey’s *post hoc*. \**p* < 0.05, \*\**p* < 0.01, \*\*\**p* < 0.001, \*\*\*\**p* < 0.0001

### 3.6 Human 5HT/glut neurons differentially express *KCNA4* in Alzheimer’s disease

We next separated tissue samples based on clinical diagnosis, comparing cognitively-normal (CN) controls to AD cases. Since we found AT8 labeling, *KCNA4* expression, and *SLC24A5* expression to be heavily biased to 5HT/glut neurons (**Figure 7A,B**), we first examined the %EA of *SLC17A8* within all 5-HT neurons. We found no difference in CN controls vs AD (**Figure 7C**, *p* = 0.2825), implying equal prevalence of 5HT/glut neurons between conditions. Similar to our findings in htau 5HT/glut neurons, we found a reduction of *KCNA4* %EA in 5HT/glut neurons (**Figure 7D**, CN 60.45 ± 6.06 % vs AD 45.73 ± 10.59 %, \**p* = 0.0437). Interestingly, we found no AD-associated difference in *KCNA4* %EA in 5HT-nonglut neurons (**Figure 7E**, *p* = 0.9024). Lastly, we assessed *SLC24A5* %EA and found no difference in 5HT/glut neurons (**Figure 7F**). We did find a reduction in *SLC24A5* %EA in 5HT-nonglut neurons (**Figure7G**, CN 4.95 ± 3.02 % vs AD 0.62 ± 0.79 %, \**p* = 0.0159), though *SLC24A5* expression levels were found to be comparatively low in 5HT-nonglut neurons overall.

**Figure 7.**
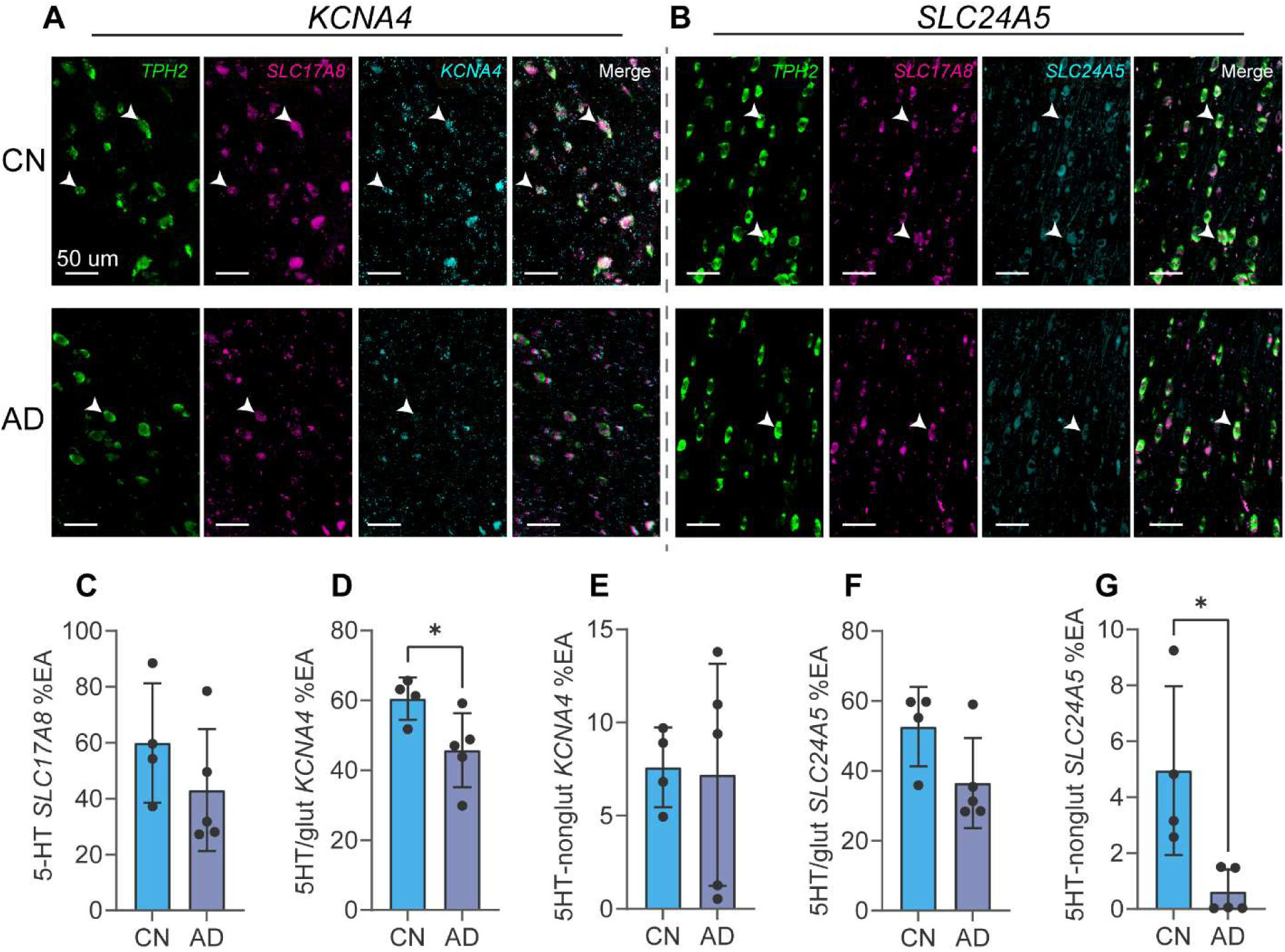
Cell type-specific reduction in *KCNA4* and *SLC24A5* in Alzheimer’s disease. ***A:*** Representative confocal images demonstrating expression patterns of *KCNA4* in 5HT/glut neurons (<u>white</u> <u>arrows</u>) for cognitively-normal (<u>CN</u>, ***top***) and Alzheimer’s disease (<u>AD</u>, ***bottom***) cases. ***B:*** Same as (***A***), but for *SLC24A5*. ***C:*** Percent expression area (<u>%EA</u>) for *SLC17A8* (VGLUT3) within *TPH2*-positive area, CN vs AD. Dataset includes all cases. ***D:*** %EA for *KCNA4* within 5HT/glut neurons, CN vs AD. ***E:*** %EA for *KCNA4* for 5HT-nonglut neurons (*TPH2*-positive, *SLC17A8*-negative). ***F,G:*** Same as (***D,E***), but for *SLC24A5*. ***C–F:*** Student’s *t*-test. ***G:*** Mann-Whitney test. \**p* < 0.05

## DISCUSSION

Using spatial transcriptomics, we identified for the first time a specific population of vulnerable 5-HT neurons within the DRN that may drive early tau pathology in both preclinical and clinical settings. Our findings demonstrate that dual-expressing 5HT/glut neurons of htau mice recapitulate the genetic and pathological features of 5HT/glut neurons in human AD, enabling the continued interrogation of AD etiology in the htau mouse model. Of these genetic factors, we find that *KCNA4* (Kv1.4) may represent a critical therapeutic target for the treatment of tau-based AD. Given the known roles of Kv1 channels in neuronal excitability, and our finding that chronic 5-HT hyperexcitability can exacerbate pTau accumulation, modulation of Kv1.4 activity and gene expression may dually contribute to the dysfunction of 5-HT downstream signaling and the trans-synaptic spread of tau pathology in AD.

The primary roles of Kv1 channels in neurons are to regulate membrane potential and modulate action potentials. These channels are highly evolutionarily conserved in *mus musculus* and *homo sapiens*, sharing 91% nucleotide and 97.2% protein sequence similarity, respectively [47]. The Agora Database, which hosts high-dimensional human transcriptomic data for Alzheimer’s disease, identifies *KCNA4* as differentially expressed in four regions thus far: reduced in the inferior frontal gyrus (log_2_FC -0.196, *p* = 0.0320), reduced in the parahippocampal gyrus (log_2_FC -0.374, *p* < 0.0001), reduced in the superior temporal gyrus (log_2_FC -0.204, *p* = 0.0287), and reduced in the temporal cortex (log_2_FC -0.379, *p* < 0.0001) [48]. Kv1.4 represents an interesting protein target that can be modified at many different levels to modulate its activity. For example, genetic ablation of a single Kv1 channel in a targeted neuronal sub-population is sufficient to drive changes in whole-animal physiology [49], and pharmacological vestibule blockers of these neuronal channels show similar effects [50,51], though no selective blocker of Kv1.4 has yet been observed or engineered. Kv1.4 channels are further endogenously modified via hetero-tetramerization [52], β-subunit modification [53], and rapid phosphorylation/dephosphorylation by PKA, PKC, and AMPK kinases [54–56]. As Kv1 channels are dampeners of excitability, reductions to their activity concomitantly increase the excitability and activity of neurons. Kv1 channels specifically regulate the repolarization phase of APs, which directly affects both the AP frequency and AP df/dt (rate of change of frequency) via effects on the AP refractory period [57]. Our electrophysiological assessments confirm these effects, with changes in the evoked 5-HT/glut excitability threshold and AP df/dt in both a current-dependent and -independent manner. Given that Kv1.4 is subject to modification at several strata beyond the gene-expression levels that we assessed, future studies that isolate its role via targeted knockout should assess protein levels and phosphorylation states of Kv1.4. Given the strong sequence similarity across Kv1 channels, any study depending on antibody specificity for Kv1.4 should be interpreted with care. In contrast to Kv1.4, less is known about the role of NCKX5 (*SLC24A5*) in neuronal function. In fact, there is a paucity of literature supporting its expression in the brain at all, with only gene-expression databases supporting its neural expression [46]. Other members of the potassium-dependent sodium/calcium exchanger family are found to be differentially expressed in various regions of the AD brain, including NCKX1, NCKX2, NCKX3, and NCKX4 [48]. Given the importance of NCKX channels in Ca^2+^ signaling [58], neural expression of *SLC24A5* warrants future investigation in the context of AD.

Next, we examined how a reduction in excitability-dampeners, similar to reduced *Kcna4* in the centromedial DRN, influences tau pathology. To do this, we employed for the first time, a chronic chemogenetics approach to investigate the relationship between increased neuronal excitability and pTau accumulation. 5HT/glut neurons project to several brain regions that mediate symptoms seen in early AD, including the hippocampus [59,60], amygdala [61,62], entorhinal cortex [63–65], and olfactory bulb and piriform cortex [65–69]. Interestingly, of the 5-HT inputs to the olfactory bulb, piriform cortex, and entorhinal cortex, the majority of these inputs may be primarily dually glutamatergic [65]. We used *Fos* expression as a proxy for the efficacy of our DREADDs effect, assuming that activation of hM3Dq would mediate increased *Fos*, and activation of hM4Di would suppress *Fos*. We adapted this approach from Zhan 2019 [70], who used Fos immunolabeling to assess chronic DREADDs activation. Our water-bottle CNO dosage was selected based on a report from Bryan Roth’s group [71], the inventor of DREADDs, who used an equivalent dose to activate hM3Dq DREADDs in the DRN for a duration of 3 weeks. Interestingly, our *Fos* RNAscope and behavior data suggest that hM4Di did not mediate chronic suppression of neuronal activity. This is interesting, given the similar EC_50_ values for CNO at hM3Dq and hM4Di reported as 6.0 nM and 8.1 nM, respectively [72,73]. Roth’s group found that anxiety-like behavioral outcomes were only elicited with short-term DREADDs activation, but not with long-term DREADDs activation, suggesting some compensatory effect within local DRN circuitry, likely involving expression of the 5-HT autoreceptors (5-HT_1A_R, 5-HT_1B_R) [74,75]. We also failed to observe robust anxiety-like behaviors after chronic manipulation, though we “washed out” the DREADDs effect before behavioral assessment to isolate any contribution of pTau accumulation to behaviors. Given that we found increased pTau accumulation in the DRN of the hM3Dq htau group, it is unsurprising that we observed a deficit in social interaction as compared to control htau mice. We previously reported similar deficits and depressive-like behaviors in htau mice as compared to C57 [23], and the data presented herein suggest that this previously-observed phenotype is at least partially driven by pTau accumulation. Unlike previous reports, we observed enhanced cognition during the NOR task in hM3Dq htau mice, despite accelerated pTau accumulation. It is tempting to speculate that chronic chemogenetic modulation of the DRN induces lasting neuroplastic changes—indeed, the hM4Di htau group also trended toward enhanced NOR performance, albeit not to a statistically significant degree. The inability of our paradigm to chronically suppress the DRN as measured via *Fos* activation, and of Roth’s paradigm to chronically modulate as measured via anxiety-like behaviors [71], support this notion that chronic modulation of the DRN somehow affects its long-term activity. Neither Fos nor 5-HT autoreceptor levels were assessed in Roth’s study, and we did not assess 5-HTR levels in this study. To extend the significance of our DREADDs findings, in the future, we will include full-cohort assessment of DREADDs activity in htau mice which includes several conditional controls and the assessment of 5-HTRs and Fos at various timepoints.

Since DREADD activation of the centromedial DRN region, which predominantly consists of 5HT/glut neurons, led to increased pTau accumulation, we next examined gene-expression and pathology patterns within the DRN of post-mortem human AD cases. We find that the anatomical organization of 5HT/glut neurons in post-mortem human DRN differs from that in the mouse DRN (i.e., no centromedial DRN). We provide evidence herein that 5HT/glut neurons in human AD are preferentially affected by tau pathology, which coincides with reduced expression of *KCNA4* in these neurons, associated with the progression of tau pathology to Alzheimer’s disease. Given that the expression levels of *KCNA4* in 5HT/glut neurons of tau-positive cases diverged based on cognitive status (**Figure 7D**), this implies that the development of tau pathology precedes dysregulation of *KCNA4*. This, in turn, suggests that dysregulation of *KCNA4* gene-expression coincides with the spread tau pathology to brain areas responsible for cognitive function. Our htau DREADDs data suggest that this may be mediated through changes in neuronal excitability that then exacerbate the accumulation and spread of tau pathology from the DRN, leading to later cognitive deficits. The primary limitation of our assessment is that we could not directly correlate the level of *KCNA4* expression with the degree of tau pathology at a single-cell level, due to the incomplete picture presented by tissue sections captured at a static timepoint. To extend the translational significance of these findings, live neuroimaging tracers for Kv1.4 should be developed.

The present study underscores the potential for spatial ‘omics technologies to change our understanding of the initiation and spread of tau pathology. By targeting the acceleration of pTau to the centromedial DRN, we were able to promote a robust depressive-like phenotype, depression being a primary characteristic of the prodromal phase of AD. These new perspectives have the power to change the landscape of AD research, enable earlier diagnosis, and may eventually yield the first cure for AD.

**AUTHOR CONTRIBUTIONS**

## ACKNOWLEDGEMENTS

We thank Mrs. Yu Xu for excellent technical assistance and help with mouse husbandry.

## DATA AVAILABILITY

The raw and processed data supporting the findings of this study are available from the corresponding author upon reasonable request.

